# Endophytic bacteria *Bacillus velezensis* NKG50 as a potential biocontrol agent of Ascochyta blight on chickpea

**DOI:** 10.1101/2025.03.20.644328

**Authors:** L. Valetti, F. Sardo, F.D. Fernández, C.S. Crociara, S. Pastor, O.A. Ruiz, M.I. Monteoliva

## Abstract

One of chickpea crops’ most devastating and economically relevant fungal diseases is *Ascochyta rabiei*, the causal agent of Ascochyta blight. The traditional agricultural management of the disease requires a complex combination of cultural, chemical, and genetic strategies. To develop a more effective management strategy, with low economic and environmental costs, we isolated 43 endophytic bacteria from asymptomatic chickpea plants native to our soil’s region. We tested their antifungal effect with mycelial growth and conidia germination inhibition tests. Our best candidate was NKG50 isolate, which inhibited *A. rabiei* growth by more than 85% in the dual test, more than 65% by cell-free supernatants, and inhibited conidia germination by 90%. This antagonistic ability was confirmed in leaflet and greenhouse assays, with a significant reduction in disease severity in both scales. The isolated NKG50 genome was completely sequenced and identified as *Bacillus velezensis* NKG50. We found 13 gene clusters associated with secondary metabolites, and five of them with an unknown function and or nature previously reported for *B. velezensis*. This study demonstrated for the first time that *B. velezensis* NKG50 is a strong candidate for the biocontrol of Ascochyta blight in Argentina.

## 1 Introduction

Chickpea (*Cicer arietinum* L.) is the third most important legume in the world, producing 18.1 million tons, distributed over an area of 14.8 million hectares (FAOSTAT, 2022). In Argentina, 90% of production is exported and the Province of Córdoba provides 50% of those exports (Carreras *et al*., 2016; Farías *et al*., 2018). Ascochyta blight, caused by the fungus *Ascochyta rabiei*, is one of the most devastating and economically important fungal diseases worldwide, capable of causing yield losses of up to 100% (Rossi *et al*., 2018). Ascochyta blight affects the leaves, stems, and pods producing lesions and breakage of buds (Pande *et al*., 2005). The disease’s development and spread occur through airborne and splashed conidia and ascospores, and infected fallow or seeds (Tivoli & Banniza, 2007). The typical agricultural management of the disease uses a combination of cultural, chemical, and genetic control. The commercial varieties most widely used in Argentina are susceptible to *A. rabiei* (Pastor *et al*., 2021, 2022). Due to the polycyclic nature of Ascochyta blight, its severity rapidly increases under congenial environmental conditions. Fungicides are not 100% effective against *A. rabiei* (Harveson & Urrea, 2013). Consequently, the disease is mitigated by repeated highly-cost fungicide applications (Banniza *et al*., 2011; Rossi *et al*., 2018). In addition to their inefficacy against Ascochyta blight, the continuous increase in the use of fungicides over time favors the emergence of pathogen resistance (Wise *et al*., 2008; Singh *et al*., 2022), and contaminates soil and water, producing an adverse effect on the quality of food and human health (Syed Ab Rahman *et al*., 2018). Therefore, it is crucial to complement and/or replace fungicides with more efficient and environmentally friendly alternatives, such as biological control agents, especially for chickpea crop in Argentina.

Endophytic bacteria are strong candidates as biological controllers since they can colonized the host, and they may benefit their host through various mechanisms, including the production of antifungal metabolites (Romero *et al*., 2021; Alenezi *et al*., 2021), siderophores (Maela & Serepa-Dlamini, 2019), hydrolytic enzymes (Myo *et al*., 2019) and inducing systemic resistance in host plants (Romero *et al*., 2019; Jacob *et al*., 2020).

Although bacterial endophytes have been extensively studied for their biocontrol effects in different crops (Muthu Narayanan *et al*., 2022), little is known about their antagonistic effect on *A. rabiei*. For instance, *Bacillus sphaericus, B. cereus*, and *B. thuringiensis* isolated from salty soils showed antifungal effects against *A. rabiei in vivo* (Rhaiem, 2020). Another study evaluated six probiotic isolates against *A. rabiei* (Moarrefzadeh *et al*., 2022). Among them, *B. subtilis* BS showed the highest mycelial growth inhibition with 57%, while *B. velezensis* JPS19 only inhibited by 47% in dual plates. Also, some mycorrhizal fungi (such as *Rhizophagus irregularis, Funneliformis mosseae*, and *Gigaspora margarita*) were tested against *A. rabiei* in chickpea varieties Bivanij and Saral in Iran, reaching up to 45% of disease suppression (Moarrefzadeh *et al*., 2021). None of them specifically isolated native endophytic bacteria from the same crop region, and the control efficiency of the pathogen was mild.

Here we aimed to isolate, classify, and identify an endophytic strain native to chickpeas grown in Argentinian soils with an antagonistic effect against the pathogen *A. rabiei* OS8 (isolated also in Argentina). Environmental factors (such as soil moisture, temperature, and carbon availability) might limit the microbiota activity for biocontrol agents once applied to the fields (Gupta et al., 2011). Then, we prioritized isolating native bacteria, more likely to survive in the Argentinian cropping region and minimized the chances of causing massive disturbances in the soil microbiota. We found the highest antifungal *in vitro* and *in vivo* effects in a *Bacillus velezensis* NKG50 strain. We sequenced the whole genome and found 13 clusters of secondary metabolites, eight previously associated with antifungal or antibacterial metabolites, and five unknown putative antibiotic metabolites and enzymes.

## 2 Materials and methods

### 2.1 Endophytic bacteria isolation and plant pathogenic fungi

Samples of healthy chickpea plants of two varieties (Kiara, Norteño, and Felipe cultivars) were collected from three field in the Córdoba Province, Argentina: site 1: (31°10’20,7’’S _ 64°00’13,3’’W) cv. Kiara; site 2 (31°03’08,4’’S _ 63°57’32,5’’W) cv. Norteño; and site 3 (31°10’20,7’’S _ 64°00’13,3’’W) cv. Felipe. Three chickpea plants in the R6 stage (60 days after planting), were randomly chosen from each field site, and the whole plants including the root system were bagged, transported to the laboratory, and stored at 4°C until further processing. The samples were collected under the permit #GOBDIGI-925099111-121 of the Dirección de Jurisdicción de Gestión de Recursos Naturales de la Secretaría de Ambiente de la Provincia de Córdoba.

Endophytic bacteria were isolated according to the method described by Valetti et al. (2018) with some modifications. Shoots, roots, and nodules were separated from each other. Each organ was carefully washed under running tap water minimizing tissue injury. Healthy, non-ruptured organs were surface sterilized by dipping in 95% ethanol for 1 min and then in 1 % NaClO solution for 3 min and rinsed five times in sterile distilled water. To confirm the endophytic nature of the isolates, 100 µl of the final rinse with sterile water was plated onto Tryptic Soy Agar (TSA, Britania) as a sterility check. One gram of sterilized leaves, roots, and nodules were macerated in a mortar utilizing 10 ml of sterile NaCl 0.9 %. The supernatant was collected, then serially diluted (10^1– 10^7) and 100 µl aliquots from the appropriate dilutions were spread on Tryptic Soy Agar (TSA) with cycloheximide 50 µg.ml^-1^ (to avoid fungal growth), per triplicate. The plates were incubated for 48 h at 28° C. All morphologically different colonies were re-streaked for the purification of the isolates. In total, 43 pure bacterial cultures were preserved at -80° C. Pathogen strain *Ascochyta rabiei* OS8 (Crociara *et al*., 2022) is stored at the Culture Collection of Instituto de Patología Vegetal (IPAVE - INTA).

### 2.2 *In vitro* screening for antagonistic activity in dual culture assay

The modified dual culture method was used to test the antagonism of the 43 isolated bacteria against *A. rabiei* (Liu *et al*., 2021). *Ascochyta rabiei* OS8 mycelia was cultivated in potato dextrose agar (PDA, Britania) plates at 21°C (12:12 h, light: dark cycle). After seven days of active growth, *A. rabiei* mycelia discs (5 mm diameter) were transferred to the fresh PDA plate. Bacteria isolates were cultured in TBS at 28° C overnight at 150 rpm. The mycelial disc was located at the center of the dish, and four drops (10 µl each) of the bacterial culture were located 2 cm from the disc and equidistant from each other. The dishes were incubated at 21° C (12:12 h, light: dark cycle). The mycelia diameters were measured daily until the control mycelia (without bacteria) reached half of the dish diameter. Percentage Inhibition of Mycelial Growth (PIRG) was calculated for each isolate according to the formula PIRG = [(control mycelia diameter – bacteria-treated mycelia diameter) / control mycelia diameter]×100 (Sharma et al. 2019).

### 2.3 Effect of extracellular metabolites in mycelial growth and conidia germination

The effect of bacteria metabolites was evaluated according to Zhang et al. (2015) with some modifications. Bacteria isolates were cultured in TBS at 28°C overnight at 150 rpm, centrifuged at 12000 g, and then, the supernatant was filtrated with a 0.22 µm pore diameter. Sterile cell-free supernatant was embedded in the PDA media (at 10 % v/v final dilution). *A. rabiei* discs were placed in the center of the Petri dish with PDA media and measured every 5 days, for 20 days. As a control, fresh TBS media was filtered and embedded in PDA media where *A. rabiei* was grown. PIRG was calculated as described above.

The same cell-free supernatants (10% v/v) were embedded in 15% (w/v) agar water for conidia germination assay. *A. rabiei* conidia suspension (500 µl of 1×10^5 conidia /ml) was spread over the solidified media and incubated at 21 °C (12:12 h, light: dark cycle) for 24 h. Conidial germination was determined by counting 100 conidia for each repetition in the magnified glasses (40X). The conidial germination percentage was calculated as each bacteria isolate supernatant count over the control count (with fresh filtrated TBS media).

### 2.4 Identification of bacterial endophytes by *16S rRNA* and *gyrB* gene sequencing

The isolates with positive cell-free supernatant antagonistic effects were taxonomically identified based on *16S rRNA* and *gyrB* gene sequencing. Total DNA was extracted using an Easy pure Genomic DNA kit (Transgene Biotech, Beijing, China). Primers used for PCR amplification were 27F (5’-AGAGTTTGATCATGGCTCAG-3), 1492R (5 -TACGGYTACCTTGTTACGACTT-3’), and gyrB-F (5-GAAGTCATCATGACCGTTCTGCAYGCNGGNGGNAARTTYGA-3’), and gyrB-R (5-AGCAGGGTACGGATGTGCGAGCCRTCNACRTCNGCRTCNGTCAT-3) (Xie *et al*., 2021). PCR was performed as follows: one cycle of 5 min at 94°C; 30 cycles of 30 s at 94°C, 30 s at 55°C, and 1.5 min at 72°C; and 10 min at 72°C. PCR products were purified and sent to Macrogen (Seoul, Republic of Korea) for Sanger sequencing. The sequences were queried against the GenBank database using the BLAST program. The phylogenetic trees were constructed using MEGA 7.0 software (Kumar *et al*., 2018).

### 2.5 Antagonistic activity in chickpea-infected leaflets

The detached leaflet assay to evaluate the *in situ* antagonistic activity of the NKG50 and HFG8 was performed according to the method of Harijati and Keane (2012). Chickpea seeds (*Cicer arietinum* cv. Kiara, susceptible to *A. rabiei*, was kindly provided by UNC-INTA) were surface-sterilized and sown in sterile agar: water 12 % (w/v) Petri dishes until germinated (∼five days). Seedlings were transferred to pots filled with sterile vermiculite and grown in a greenhouse (21 °C, 12:12 h light: dark cycle). The youngest fully expanded leaves were collected from 30-day-old plants. Detached leaflets were surface sterilized by sequential immersion in 70% (v/v) ethanol 30 s, and 1% (w/v) NaOCl 4 min. They were rinsed five times with sterile water and immersed for 15 min in the saturated bacterial culture grown overnight (1×10^8 ufc/ml), or in the sterile filtered culture (as previously described). Then, the leaflets were placed in Petri dishes containing sterile agar water 10%.

Infections with the pathogen (*A. rabiei* OS8) were carried out according to Bayraktar et al. (2016). The spores were obtained from a 14-day culture grown in chickpea seed meal agar (CSMA: 40 g chickpea flour, 20 g sucrose, 12 g agar per liter of water) at 21 °C (12:12 h, light: dark cycle). The spore suspension was adjusted to a concentration of 1×10^5 spores/ml in sterile water. Each leaflet was inoculated with 5 µl of spore suspension and kept for 5 days at 21°C (12:12 h, light: dark cycle). Three Petri dishes were used for each treatment and each dish contained a leaf with 12-14 leaflets. Lesions were evaluated with transillumination. Disease incidence (%) was expressed as the proportion of symptomatic leaflets over the total. The damaged area percentage was estimated using ImageJ software over the total leaflet area. The experiment was replicated two times with three replicates.

### 2.6 Greenhouse experiments

The greenhouse experiments were carried out according to the protocol of Crociara et al. (2022) with modifications, in the winter of 2021 (one replicate) and 2024 (two replicates). Chickpea (*Cicer arietinum* cv. Kiara) seeds and Ascochyta spores were prepared as described above. After seven days, seedlings were transferred to sterile vermiculite-filled pots of 360 cm^3^, in a greenhouse under semi-controlled conditions (21 °C, 12:12 h, light: dark cycle). The bacterial cultures of selected strains (NKG50 and HFG8) were applied by foliar spray (2 ml per plant) at the time of infection (1×10^8 cfu/ml). Two-week-old plants (at V4) were infected with 4 ml of spore suspension (1×10^5 spores/ ml), using a hand atomizer (Valetti *et al*., 2021). After the infection, the plants were kept at a relative humidity (RH) of 100% for 48 h, and of 65% RH for the rest of the experiment (optimum conditions for the *A. rabiei* infection, Pande et al., 2011). Disease severity was evaluated 14 days after inoculation using a 0– 9 rating scale, as follows: 1 = no visible lesions, 3 = only flecks on leaves, 5 = lesions on stems and flecks on leaves, 7 = stem breakage in injured areas, and 9 = dead plant (Nasir et al., 2000, modified from Singh et al., 1981). Infected pots without the bacteria were used as controls (sick check). The experimental unit consisted of one plant per pot and 10 pots were assigned per treatment, distributed randomly, and repeated in two different seasons (three times in total). The severity was determined as the average of the 10 pots.

### 2.7 NKG50 genomic DNA purification and sequencing

Genomic DNA (gDNA) was isolated from pure cultures using a commercial kit (Wizard Genomic DNA Purification Kit, Promega) and quantified using Nanodrop® (N-1000) and Quantus® (Promega, USA). Sequencing libraries were constructed using the Rapid Barconding Kit (SQK-RBK004) for ONT platform sequencing in a MinION Mk1B device and flowcell R.9.4.1 chemistry. ONT RAW signals were base-called using guppy v 6.4.2 in the HAC model (model dna_r9.4.1_450bps_hac.cfg).

### 2.8 *B. velezensis* NKG50 genome assembly and annotation

The ONT reads were filtered using Filtlong (https://github.com/rrwick/Filtlong) by removing reads with a length of less than 1000 bp and those with an average quality score of less than 8. The assembly was conducted using Flye assembler (https://github.com/fenderglass/flye) using the -nano-raw mode. The final assembly was polished by applying two rounds of medaka (https://github.com/nanoporetech/medaka). The completeness of the final polished assembly was confirmed using BUSCO (https://github.com/WenchaoLin/BUSCO-Mod). The annotation process was conducted using the NCBI Prokaryotic Genome Annotation Pipeline (Tatusova et al., 2016). KAAS-KEGG Automatic Annotation Server (https://www.genome.jp/kegg/kaas/) was used for the functional characterization of protein-coding regions and reconstruction of metabolic pathways. A graphical map of the complete genome was resolved using the GenoVi tool (Cumsille *et al*., 2023).

### 2.9 *B. velezensis* NKG50 orthologues clustering and phylogenetic analyses

Orthologous protein clusters were identified using Orthofinder v2.5.2 (https://github.com/davidemms/OrthoFinder). The genome sequences of representative *Bacillus species* were obtained from GenBank (Table S1). For phylogenetic analyses, alignments of concatenated nucleotide sequences of single-copy core genes were constructed using MAFFT v7.450 within Geneious R.10 software (Biomatters Ltd., Auckland, New Zealand). Phylogenetic trees were inferred with IQ-TREE (http://www.iqtree.org/) using an automatic substitution model and ultrafast bootstrap analysis with 1000 replicates. Whole-genome comparisons were conducted using fastANI v1.1 (Jain *et al*., 2018). We also computed the genome-to-genome distance calculator online tool (https://ggdc.dsmz.de/home.php) to calculate dDDH values against *Bacillus velezensis* strain NKG50 sequenced in this study.

### 2.10 Statistical analysis

The statistical analysis was performed using the Infostat 2020 statistical package (Di Rienzo *et al*., 2020). Data were subjected to Analysis of Variance (ANOVA), followed by mean comparison by multiple-comparisons method (DGC) in *in vitro* experiments and Least Significant Difference (LSD), with a 5 % significance level for leaflet assay and pot trial.

## 3 Results

### 3.1 Screening of chickpea bacteria and cell-free supernatants with antagonist activity against *Ascochyta rabiei*

A total of 43 isolates were obtained from leaves, roots, and nodules of asymptomatic chickpea plants, cultured in fields of Cordoba Province (Argentina). Their antagonistic capacity against the *Ascochyta rabiei* OS8 strain was tested in dual plates. Among all, 16 isolates showed inhibitory activity higher than 50% (Fig. 1, Fig S1).

**Figure 1.**
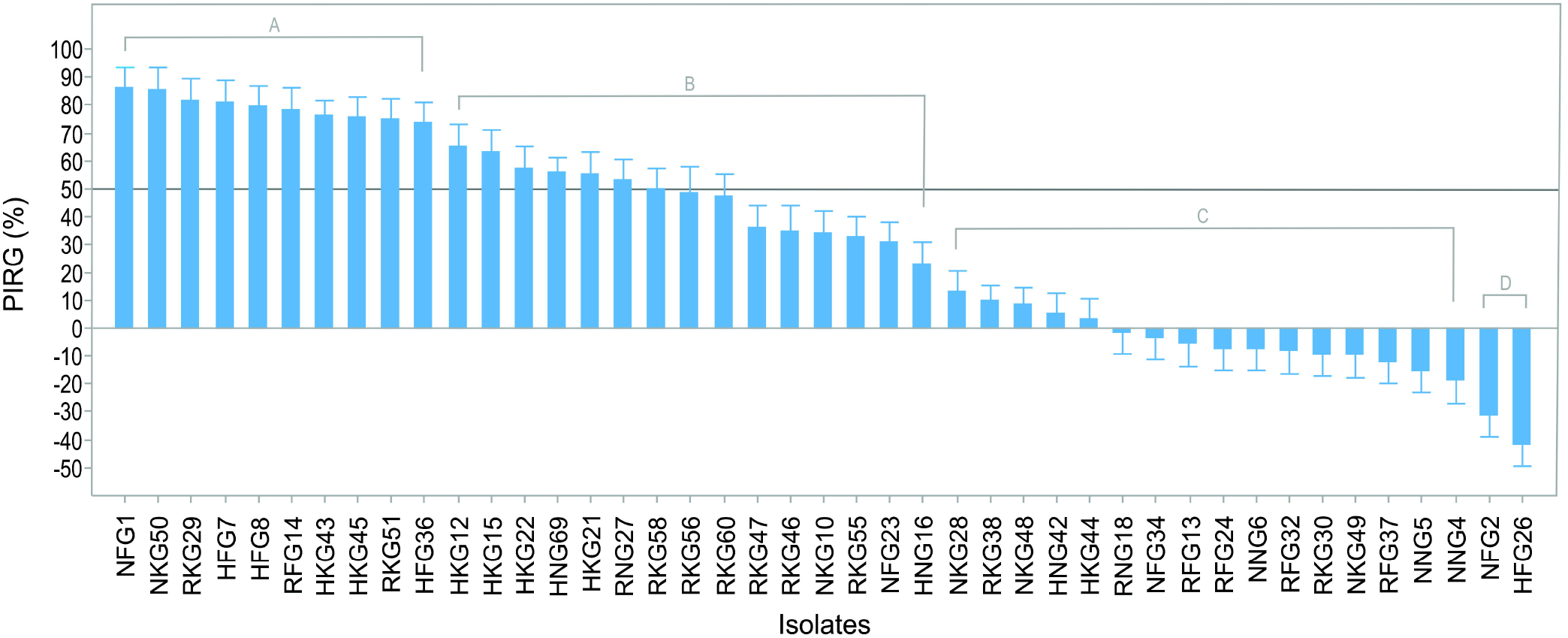
Antifungal effect of bacterial endophytes from chickpea plants against *Ascochyta rabiei*. Dual plate screening of 43 bacteria isolates showing percentage inhibition of mycelial growth (PIRG). Data represent the mean ± SE of two independent replicates with five repetitions each. Shared letters indicate no significant difference (p>0.05, ANOVA, DGC).

Sterile cell-free supernatants of those 16 isolates were tested for their capacity to inhibit mycelial growth and conidia germination (as the antagonist effect of the secreted metabolites). Eight isolate’s supernatants inhibited growth by ≥60% (Fig. 2A, bars). Only three isolates highly inhibited conidia germination (HFG8, HKG21, and NKG50 by ≥90%) (Fig. 2A, squares). Although HKG21 showed a high conidia inhibition, only showed a limited mycelial growth inhibition by their metabolites. Then we prioritize to continue studying NKG50 and HFG8 (Fig. 2B, C).

**Figure 2.**
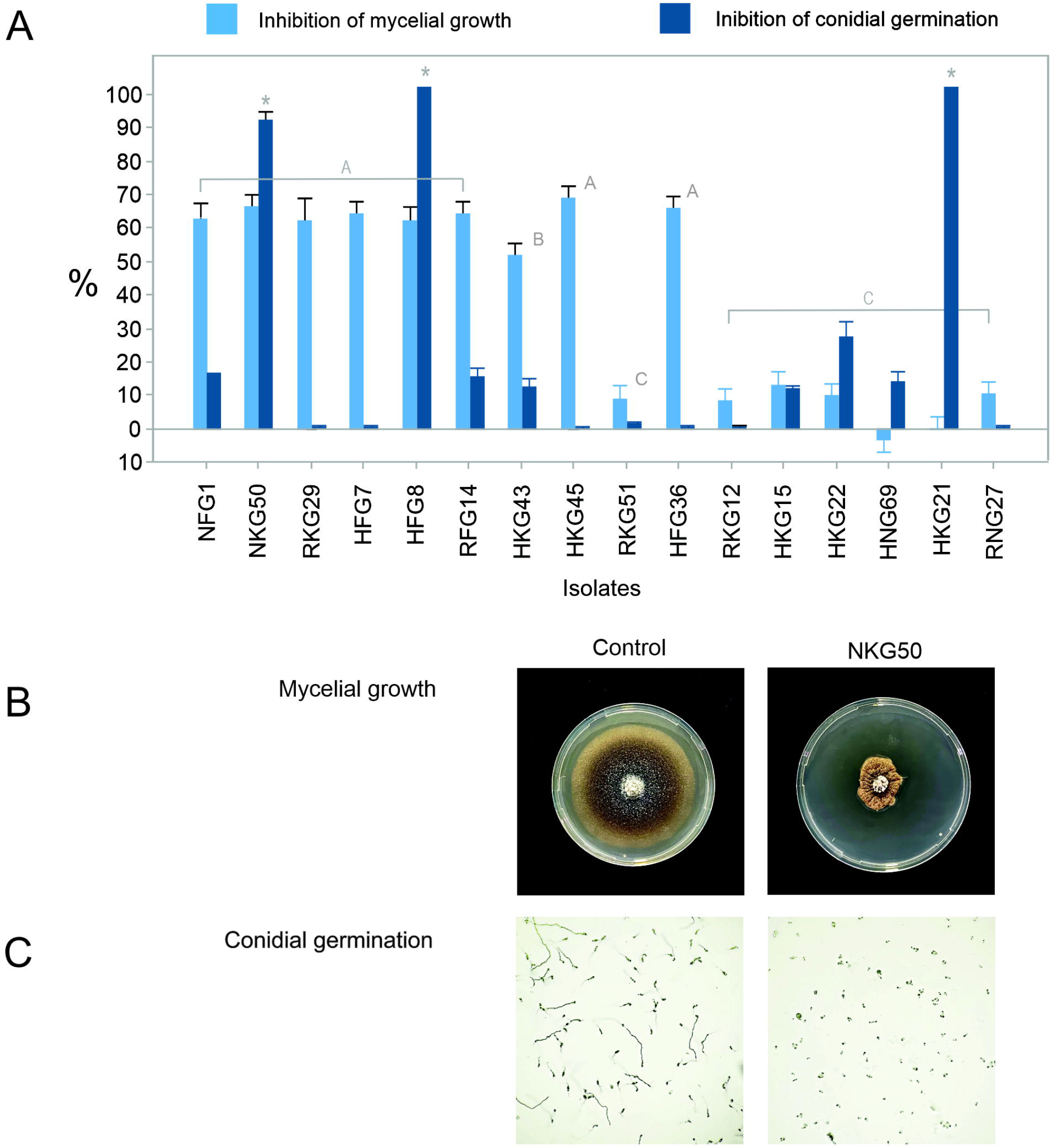
Evaluation of the cell-free supernatant effect on *A. rabiei* growth and conidia germination. **(A)** *A. rabiei* mycelial growth (light blue) and conidia germination rate (dark blue) in the presence of the cell-free supernatant isolates; **(B)** Fungal colony growing in the absence (Control) or presence (NKG50) of the cell-free supernatant of NKG50 isolate, and **(C)** pictures of *A. rabiei* mycelia and conidia incubated at 21 °C for 18 h in the absence (Control) or presence (NKG50) of the cell-free supernatant of NKG50 isolate. In A, data represent the mean ± SE of two independent replicates with five repetitions. Shared letters above mycelial growth inhibition bars indicate no significant difference (p>0.05, ANOVA, DGC). In B, asterisks (*) above the conidia germination rate bars indicate significant differences (p<0.05, ANOVA, DGC).

### 3.2 Bacterial identification

According to their antifungal effect in dual plates and cell-free supernatant, the best ten isolates were sequenced, using the *16S rRNA* and *gyrB* genes. All isolates showed a high percentual identity with the *Bacillus* genus for both regions (Table S1). The phylogenetic analysis revealed that NKG50, HFG7, HKG45, NFG1, and RFG14 clustered together with *Bacillus velezensis/ amyloliquefaciens* operational group (Fan *et al*., 2018), with >97% sequence similarity for the *16S rRNA* gene sequence (Fig. 3C), and >98% sequence similarity for the *gyrB* gene (Fig. 3D). HFG8 and HFG36 clustered with *B. subtilis*, while HKG43 and RKG29 clustered differently with *16S* and *gyrB* genes.

**Figure 3.**
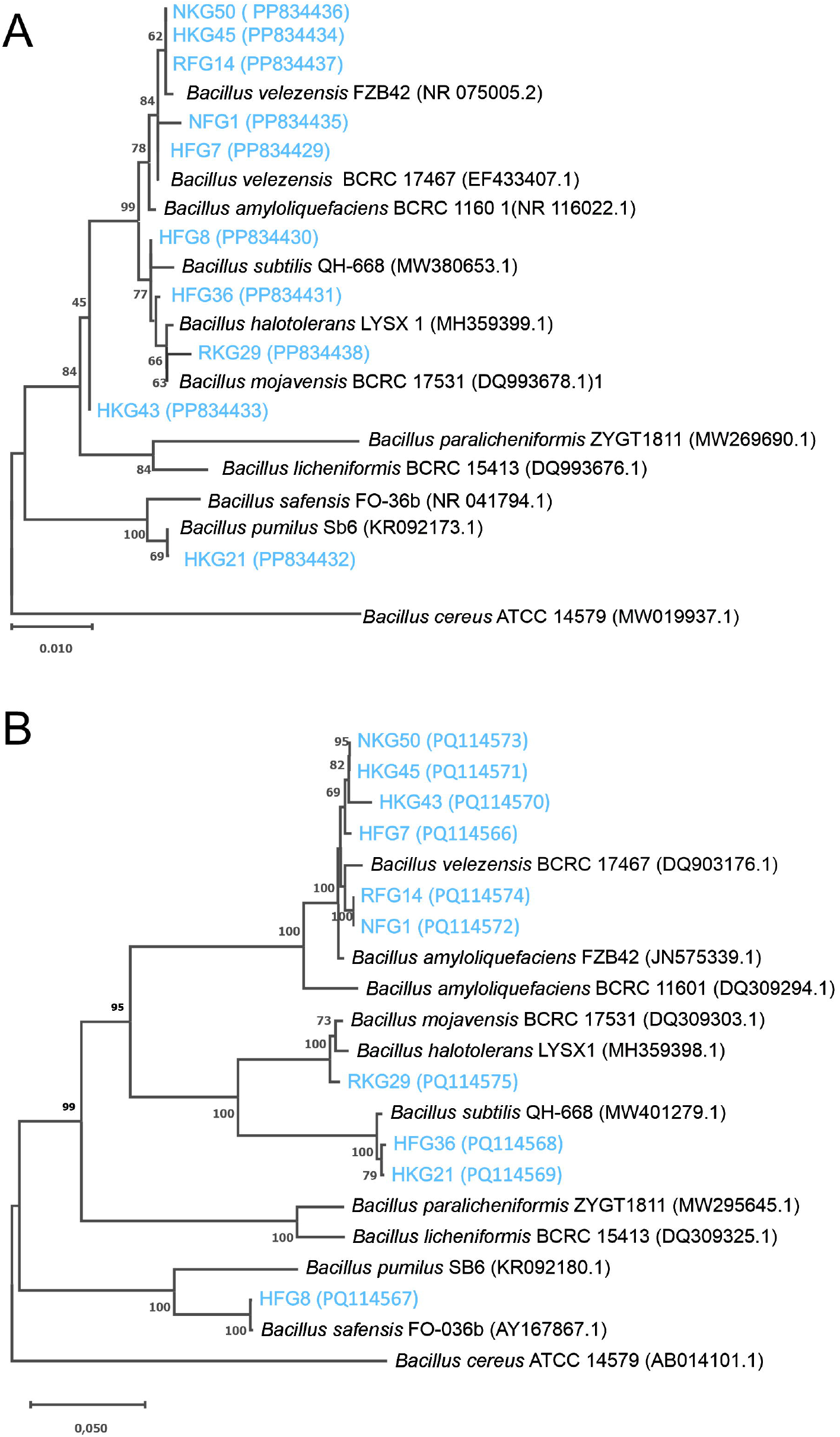
Phylogenetic analysis of the *16S rRNA* (A) and *gyrB* (B) genes. The 10 isolates with the highest antifungal effect of cell-free supernatant assay against *A. rabiei* mycelial growth and conidial germination were analyzed with the closest species within the genus *Bacillus*. The phylogenetic trees were constructed using the neighbor-joining method. The number at each node represents the percentage of times the group of strains in that branch occurred based on 1000 bootstrap cycles. The scale bar indicated 0.05 substitutions per nucleotide position.

### 3.3 Antifungal effect in chickpea plants infected with *A. rabiei*

Given that NKG50 and HFG8 strains showed the highest inhibition of mycelial growth (in dual plates and cell-free supernatants) and conidia germination, we evaluated the antifungal effect of these two isolates in two different approaches: *in situ* (leaflets) and *in vivo* (whole plants).

First, detached chickpea leaves were treated with each isolate culture or their cell-free - sterile-supernatant and then infected with *A. rabiei* conidia suspension. Symptoms were evaluated on the fifth day after infection. The inoculation with NKG50 strain reduced the incidence and damaged area by more than 90%, while the cell-free supernatant reduced the damaged area by almost 60% but not the incidence (Fig. 4). The inoculation with HFG8 cell culture reduced the incidence by less than 40% and the damaged area by less than 65% compared with the sick control. HFG8 cell-free supernatant had no protective effect.

**Figure 4.**
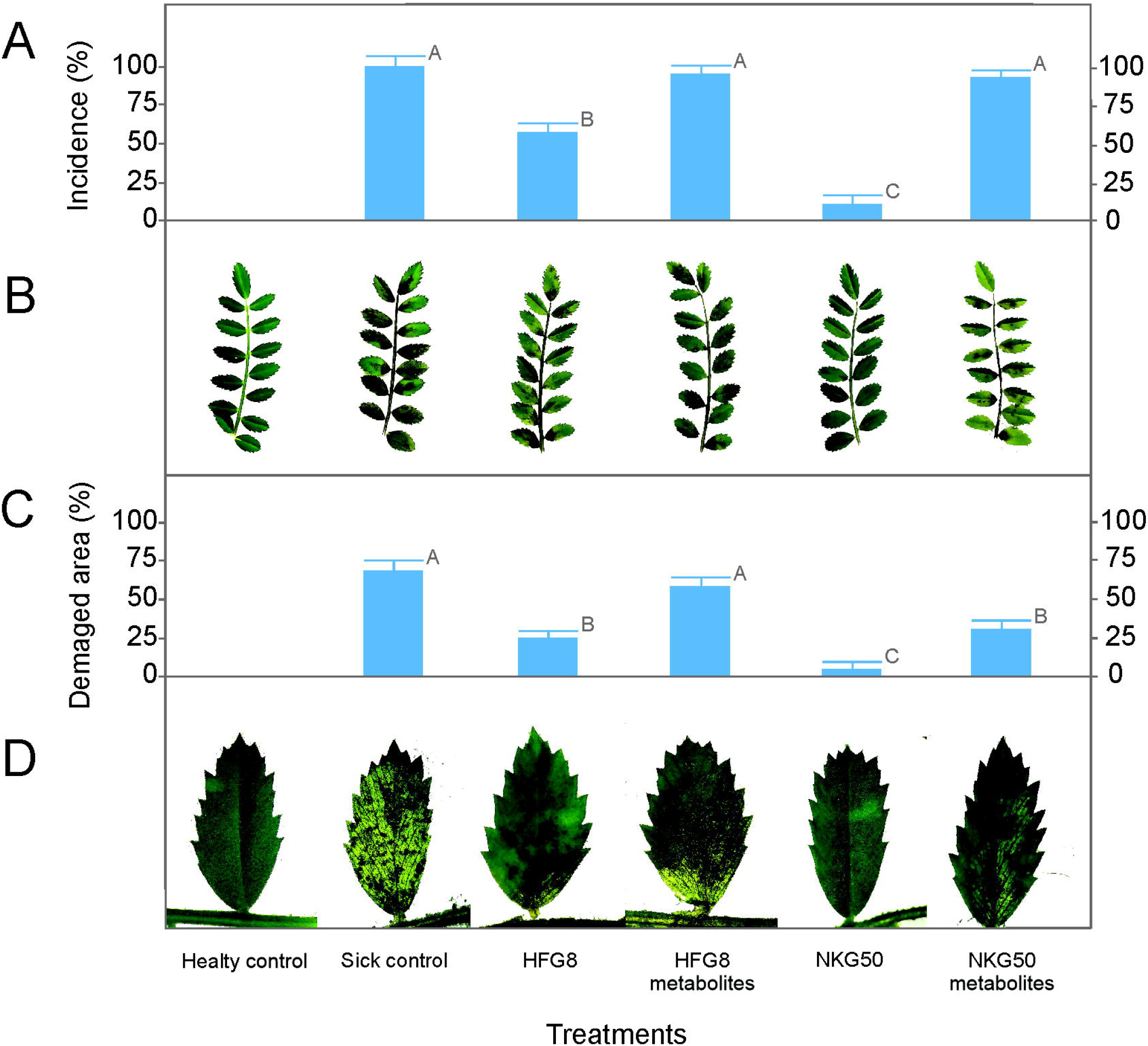
Antifungal effect of NKG50 and HFG8 strains on infections with *A. rabiei* in chickpea leaflet assay. **(A)** Incidence as a percentage of symptomatic leaflets per leaf, after inoculation with culture or cell-free supernatant of NKG50 or HFG8, and infected with *A. rabiei* one day after the inoculation. **(B)** Pictures of representative leaf quantified in A. **(C)** Damage area percentage of symptomatic leaflets averaged per leaf, after inoculation with culture or cell-free supernatant of NKG50 or HFG8, and infected with *A. rabiei* one day after the inoculation. **(D)** Pictures of representative leaflets quantified in C. Lesions were evaluated with transillumination. In A and C, data represents the mean ± SE of two independent replicates with five repetitions. Shared letters indicate no significant difference (p>0.05, ANOVA, LSD).

Secondly, we validated the biocontrol activity of the isolate NKG50 in greenhouse experiments (2021 and 2024). The whole plants were sprayed with NKG50 culture media, and 48 h were infected with *A. rabiei* conidia suspension. NKG50 significantly reduced severity, from 7 to 4 grade compared to the sick plants (Fig. 5). In all infected plants, treated or not with the bacteria, the incidence was >90% (not shown). The cell-free supernatants did not affect disease severity or incidence (not shown).

**Figure 5.**
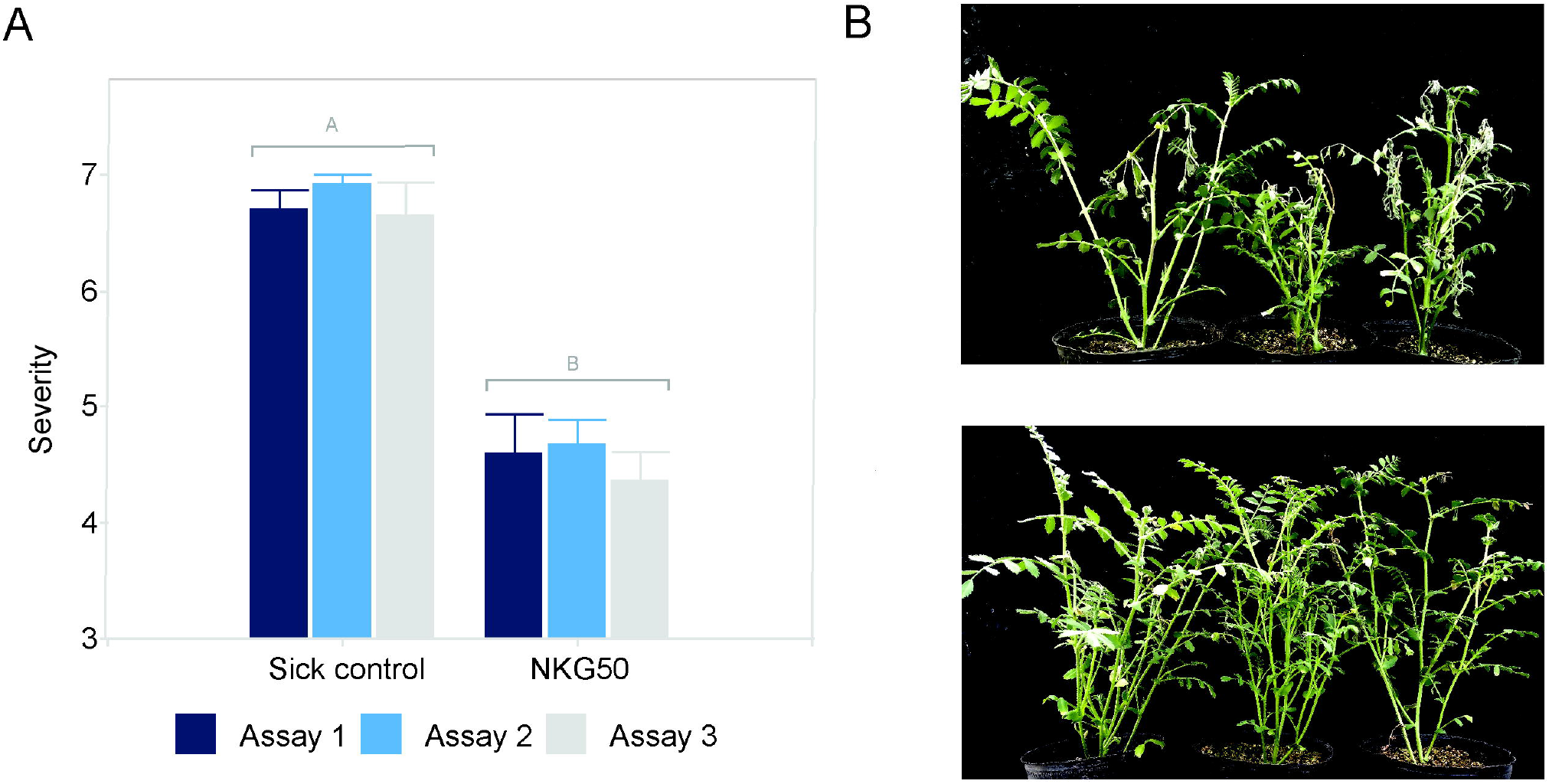
Antifungal effect of NKG50 strain on *A. rabiei* infections of chickpea plants. The bacterial suspension was applied by foliar spraying in vegetative stage V4, before the *A. rabiei* infection. **(A)** Disease severity in chickpea plants treated or not with the B. velezensis NKG50 cell culture; **(B)** Picture plants infected with *A. rabiei* (Sick control, top panel), or plants pre-treated with NKG50 strain before infection with *A. rabiei* (NKG50, bottom panel). In A, data represents the mean ± SE of three independent replicates with ten repetitions. Different letters indicate a significant difference (p>0.05, ANOVA, LSD).

### 3.4 Assembly and key features of *Bacillus velezensis* NKG50 genome

Given the effective antifungal effect of the NKG50 strain, we sequenced the whole genome. 1.3 Gb of ONT reads (72.589 reads, N50 19 kb) were generated (GenBank SRA PRJNA1123166). The final genome was assembled into a single circular contig, comprising 4.123.916 bp (46.00% GC) (Fig. 6, Table S1, Table S2). In the annotation process, 4.062 predicted CDSs (including 458 hypothetical proteins), nine complete rRNA operons (5S, 16S, 23S), and 85 tRNAs were annotated in the chromosome (Table S2). Based on BUSCO analyses, the assembly was found to be 94.4% complete (with 425 complete and single-copy BUSCO groups out of 450 total BUSCO groups from the bacilalles_odb10 lineage). That assembly level covered most of the coding regions and confirmed the quality of the genome assembly.

**Figure 6.**
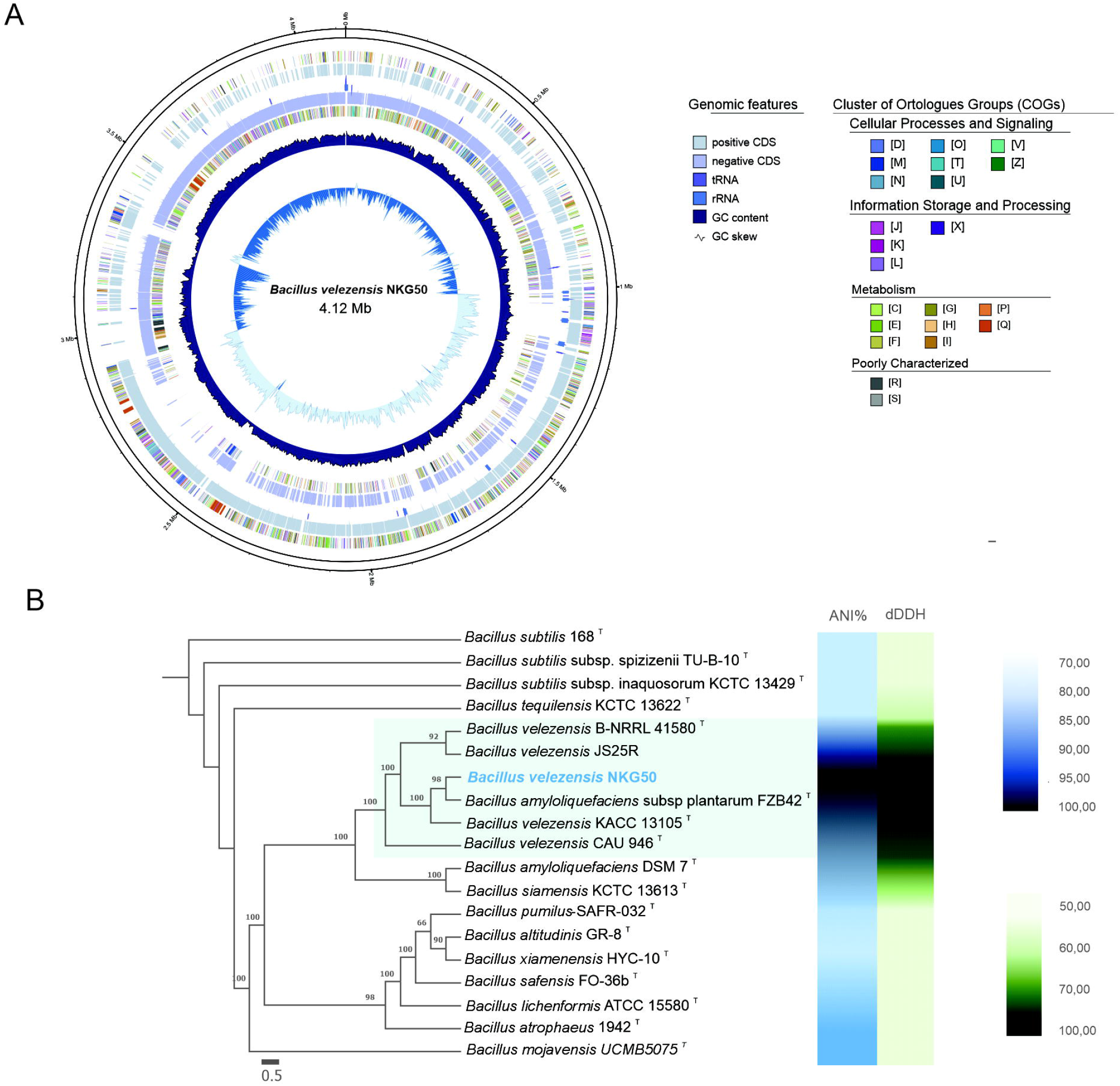
(A) Circular map representation of the complete genome of *Bacillus velezensis* NKG50. Labeling from outside to inside: forward contigs (first circle), forward COGs (2nd), forward CDS and tRNAs (3rd), forward RNAs (4th), reverse CDS and tRNAs (5th), reverse RNAs (6th), reverse COGs (7th), GC content (8th) and GC skew (9th). **(B)** Maximum likelihood phylogenies. The numbers on the internal branches indicate the level of bootstrap support based on 1,000 resampling; only values ≥60% are shown. The concatenated alignment contains 58 genes (Single-copy coding genes shared by all strains) and 40.806 aligned nucleotide sites. In light blue B. velezensis co-species clade. Bar: 0.5 substitutions per nucleotide position. Genome-to-genome comparison statistics of *B. velezensis* NKG50 among all genomes analyzed in this study. Average nucleotide identity (ANI) values were calculated using fast ANI v1.1, and dDDH values were determined using the genome-to-genome distance calculator online tool (https://ggdc.dsmz.de/home.php). The scales for ANI% (blue) and dDDH (green) are shown on the right.

### 3.2 3.5 Phylogenetic analysis of the *Bacillus velezensis* NKG50 whole genome

For phylogenomic evaluation, the *B. velezensis* NKG50 was compared to 18 other selected *Bacillus* species (five belonging to the *Bacillus velezensis* clade, Table S1). Orthologue analysis revealed the presence of 58 single-copy genes (SCGs) common to all species. The phylogenetic tree, based on DNA sequence from the alignment of these 58 SCGs (40.806 bp), confirmed that the *B. velezensis* NKG50 genome, previously identified by sequencing *16S* and *gyrB* genes, clustered within the other *B. velezensis* genome species and form a monophyletic group with high support value (100) (Fig. 6A).

To obtain genome distance metrics and confirm the taxonomic identification, all 18 genome sequences were evaluated by the average nucleotide identity (ANI) and digital DNA-DNA hybridization (dDDH). The *B. v*elezensis NKG50 genome showed >96% in ANI comparison and >70% for dDDH (Richter & Rosselló-Móra, 2009) (Fig. 6B, Table S2), confirming the taxonomic identification as *B. velezensis* by the phylogenomic analysis.

In addition, we explored the genome of *B. velezensis* NKG50 looking for secondary metabolite gene clusters, and identified 13 of them. Eight showed significant similarities with the previously characterized clusters of secondary metabolites, such as bacillibactin, subtilin, bacilycin, surfactin, macrolatin H, bacillaene, fengicyn, and dificidyn (all with 100% identity), while one exhibited a low value (butirosin A/ butirosin B, with 7%). Four out of the 13 clusters codify unknown putative antibiotic metabolites (such as terpenes and lanthipeptides), and polyketide enzymes, in the antiSMASH database (Table 1).

## 4 Discussion

The development of biological control for agricultural inputs using antagonistic bacteria is one of the most promising alternatives to chemical fungicides. Several species of non-pathogenic bacteria, mainly *Bacillus, Pseudomonas*, and *Paenibacillus* genera, have been used as potential biocontrol agents against soil-borne pathogens. We tested the antagonistic and biocontrol capacity *in vitro* and *in vivo* of 43 isolates native to the Argentinian cropping region. All the most effective antifungal isolates belong to the genus *Bacillus* (Fig. 2, Table S1). *Bacillus* species have attracted the attention of researchers and industries because their long-term viability facilitates the development of commercial products (Qiao *et al*., 2014). They also produce a broad spectrum of antimicrobial compounds and fungal cell-wall degrading enzymes (Myo *et al*., 2019).

Among all the isolates, the *B. velezensis* NKG50 from chickpea nodules was the most effective in reducing *A. rabiei* growth and conidia germination. *B. velezensis* species was first isolated in 1998 and sequenced in 2007 (Ruiz-García *et al*., 2005; Chen *et al*., 2007; Fan *et al*., 2018). Since then, it has been extensively studied for its plant growth promotion and biocontrol effects against several fungal pathogens such as *Ralstonia solanacearum* and *Fusarium oxysporum* (Cao *et al*., 2018), *Colletotrichum gloeosporioides* (Jin *et al*., 2020), or *Botrytis cinerea* (Stoll *et al*., 2021), among others. Given that Ascochyta blight is a polycyclic disease, with possible devasting spread and crop damage, *B. velezensis* NKG50 has the potential to reduce not only the disease severity by inhibiting the pathogen growth but also to reduce the secondary infections, by inhibiting conidia germination.

Here we sequenced the whole genome and confirmed the identity of our isolate NKG50 as a *B. velezensis*. By genome mining, we find the 13 secondary metabolite clusters usually reported for other *B. velezensis* strains (Fan *et al*., 2018; Wang *et al*., 2019; Shen *et al*., 2023; González-León *et al*., 2024). According to that prediction, *B. velezensis* NKG50 has the potential to produce several antimicrobial metabolites from different groups: lipopeptides (iturin, fengycin, and surfactin), polyketides (macrolactin, bacillaene, and difficidyn), and peptides (bacilysin). Some of them might be responsible for the fungal inhibition shown here. Lipopeptides produced by *B. velezensis*, such as iturin and fengycin, are the major inhibitors of *Fusarium oxysporum* and *Colletotrichum gloeosporioides* (Cao *et al*., 2018; Jin *et al*., 2020). However, the antifungal effect caused by *B. velezensis* inoculation is mediated by intertwined effects. Several antifungal metabolites have been reported to have direct and indirect effects. For instance, fengycin and bacillomycin have antibiotic activity but also indirectly induce plant defense responses (Fan *et al*., 2018), which hinder the effects of some specific metabolites. Further studies on *B. velezensis* NKG50 are underway to identify and demonstrate the secondary metabolites responsible for the antifungal effect against *A. rabiei*, and its ability to induce ISR.

Lastly, a few differences could be detected by comparing the clusters of secondary metabolites in different *B. velezensis* strains. For instance, *B. velezensis* NKG50 did not seem to have the ability to produce mersacidin, a lanthipeptide class II, predicted in *B. velezensis* SH-1471 (Shen *et al*., 2023), or bacteriocin, a non-ribosomal peptide predicted in *B. velezensis* GUMT319 (Ding *et al*., 2021). However, it does have the clusters coding subtilin which none of those strains has on their genome. These differences among genomes might enable different antifungal effects against pathogenic fungi in *B. velezensis* isolates, which has not been addressed yet.

We isolated the native *Bacillus velezensis* NKG50 strain from chickpea nodules with high potential as a biocontroller agent of *Aschochyta rabiei*. NKG50 strain strongly reduced conidia germination and mycelial growth. We successfully sequenced the complete genome of *B. velezensis* NKG50, providing additional insights into the gene repertoire associated with antibiotic effects, which are crucial for understanding its behavior as a biocontrol agent. Furthermore, the genomic predictions point out that *B. velezensis* NKG50 has the potential to control several other agronomically relevant pathogens.

## Author Contributions

LV: Conceptualization, formal analysis, funding acquisition, investigation, methodology, project administration, supervision, visualization, writing original draft and review; FS: research, writing – review & editing; FDF: investigation, research, software, visualization, writing – original draft; CSC: research, resources, writing – review & editing; SP: research, resources, funding acquisition, writing – review & editing; OAR: supervision, funding acquisition, writing – review & editing; MIM: Conceptualization, funding acquisition, investigation, methodology, project administration, visualization, writing original draft, editing & review.

## Acknowledgments

FS and CSC are CONICET fellows. OAR and MIM Career Investigators of CONICET. LV, FDF, SP, and MIM, are Investigators of INTA. The authors thank Agustin Carbajal for the critical reading of the manuscript.

## Data Availability Statement

The genomic datasets generated for this study can be found in the NCBI Sequence Read Archive (SRA) under the accession SRX24891422. The complete genome of *Bacillus velezensis* strain NKG50 was deposited in GenBank under the accession CP158307.1 (BioProject: PRJNA1123166, BioSample: SAMN41798565). The rest of the datasets supporting the conclusions of this article will be made available by the authors on request, without undue reservation.

---

**Figure S1. Dual plates assays between *Ascochyta rabiei* mycelia and the 43 bacterial isolates, after 21 days of incubation at 21 °C.** Representative pictures of the isolates plotted in Figure 1 were chosen.

