## Supplementary figures and images for "Endophytic bacteria *Bacillus velezensis* NKG50 as a potential biocontrol agent of Ascochyta blight on chickpea"

### Fig S1

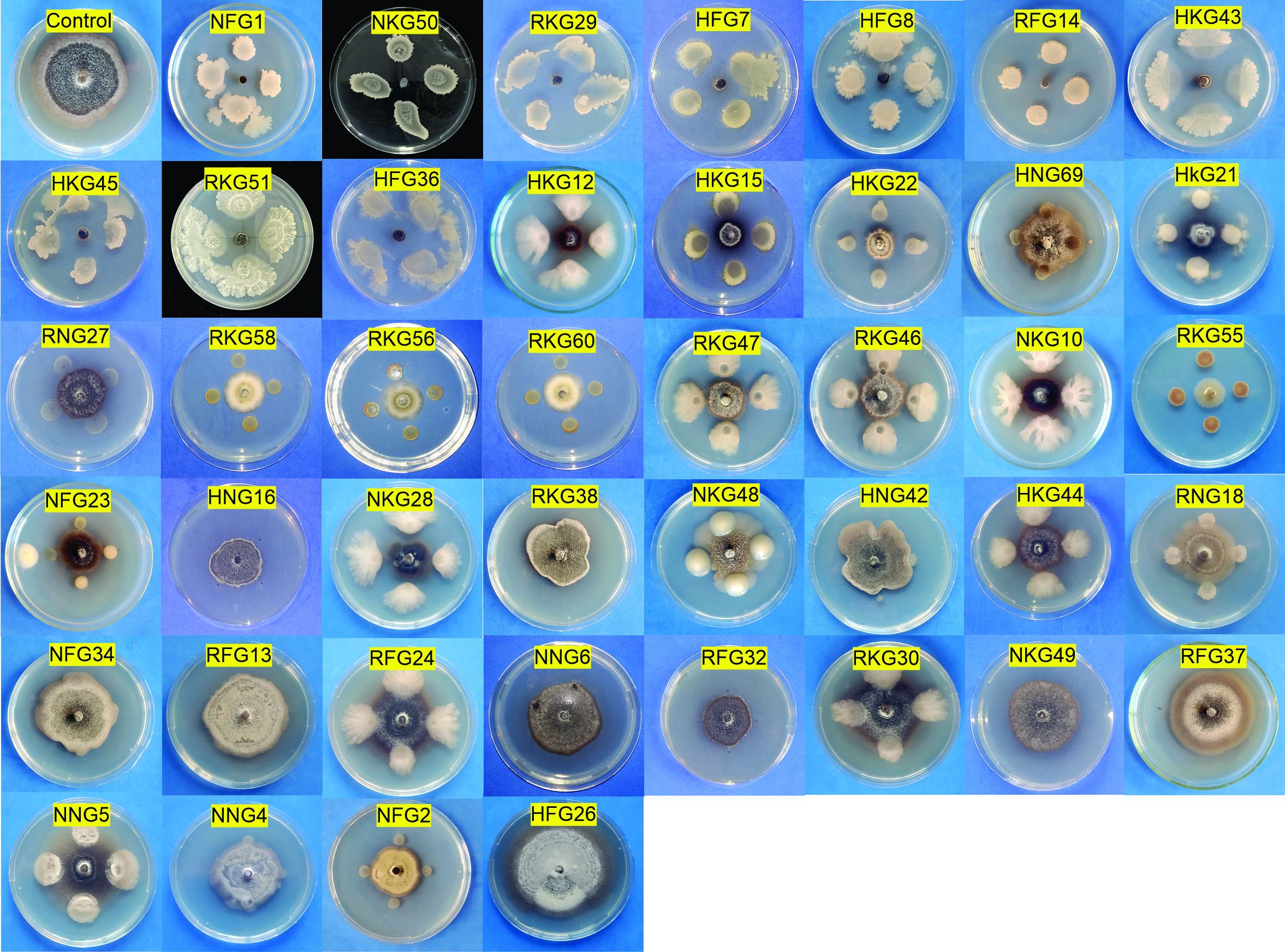
