## Supplementary material for "Endophytic bacteria *Bacillus velezensis* NKG50 as a potential biocontrol agent of Ascochyta blight on chickpea": Table 1

**Table 1. Potential gene clusters involved in synthesizing secondary metabolites in *B. velezensis* NKG50.** "From" and "To" indicate the predicted cluster's beginning and end (in base pairs). Abbreviations: nd, no data; NRPS, non-ribosomal peptide synthase; PKS, polyketide synthase; RiPP, ribosomal synthesized and post-translationally modified peptide.

| Cluster | Most similar known compound | Function | Biosynthetic class | Similarity | From | To | bp | Reference |
|---|---|---|---|---|---|---|---|---|
| 1 | Bacillibactin | siderophore, antibacterial | NRP-metallophore, NRPS, RiPP-like | 100% | 80,644 | 131,163 | 50,519 | Chen et al., 2008; Dimopoulou et al., 2021 |
| 2 | Subtilin | antibacterial | lanthipeptide-class-I, RiPP | 100% | 282,354 | 308,922 | 26,568 | Shettar et al., 2023 |
| 3 | Bacilysin | antifungal, antibacterial | other | 100% | 653,090 | 694,508 | 41,418 | Wu et al., 2015 |
| 4 | Surfactin | antifungal | NRP: lipopeptide | 91% | 1,314,246 | 1,379,654 | 65,408 | Koumoutsi et al., 2004 |
| 5 | nd | nd | lanthipeptide-class-V | nd | 1,791,773 | 1,833,630 | 41,857 | |
| 6 | Butirosin A/ B | antibacterial | PKS-like, saccharide | 7% | 1,949,012 | 1,990,256 | 41,244 | Howells et al., 1972 |
| 7 | nd | nd | terpene | nd | 2,073,037 | 2,093,777 | 20,740 | - |
| 8 | Macrolactin H | antifungal | transAT-PKS, polyketide | 100% | 2,372,396 | 2,460,622 | 88,226 | Liu et al., 2016 |
| 9 | Bacillaene | antibacterial / ISR induction | transAT-PKS, T3-PKS, NRPS-like, polyketide+NRP | 100% | 2,724,180 | 2,834,295 | 110,115 | Chen et al., 2006 |
| 10 | Fengycin | antifungal/ ISR induction | NRPS, transAT-PKS, NRPS-like, betalactone | 100% | 2,902,137 | 3,039,951 | 137,814 | Chen et al., 2007 |
| 11 | nd | nd | terpene | nd | 3,106,859 | 3,128,742 | 21,883 | - |
| 12 | nd | nd | T3-PKS | nd | 3,194,156 | 3,235,256 | 41,100 | - |
| 13 | Difficidin | antibacterial | transAT-PKS | 100% | 3,488,147 | 3,594,317 | 106,170 | Koumoutsi et al., 2004 |
