## Supplementary material for "Endophytic bacteria *Bacillus velezensis* NKG50 as a potential biocontrol agent of Ascochyta blight on chickpea": Table S1

**Table S1. Isolates identification in the NCBI database by sequencing *16S* and *gyrB* genes**

| Gene | Strain | Species with the highest similarity | Identity (%) | Cover (%) | GenBank number |
|---|---|---|---|---|---|
| ***16S*** | HFG8 | *Bacillus subtilis* DSM 10 | 100 | 97 | NR_027552.1 |
| | NKG50 | *Bacillus velezensis* FZB42 | 99.8 | 98 | NR_075005.2 |
| | HFG7 | *Bacillus velezensis* FZB42 | 99.6 | 96 | NR_075005.2 |
| | HFG36 | *Bacillus subtilis* DSM 10 | 99.9 | 98 | NR_027552.1 |
| | HKG21 | *Bacillus safensis* NBRC 100820 | 100 | 100 | NR_113945.1 |
| | HKG43 | *Bacillus subtilis* subsp. *inaquosorum* BGSC 3A28 | 99.27 | 98 | NR_104873.1 |
| | HKG45 | *Bacillus velezensis* FZB42 | 99.6 | 98 | NR_075005.2 |
| | NFG1 | *Bacillus velezensis* FZB42 | 97.4 | 98 | NR_075005.2 |
| | RFG14 | *Bacillus velezensis* FZB42 | 99.4 | 99 | NR_075005.2 |
| | RKG29 | *Bacillus mojavensis* IFO15718 | 99.7 | 99 | NR_024693.1 |
| ***GyrB*** | HFG8 | *Bacillus subtilis* subsp. *subtilis* 168 | 97.5 | 100 | CP053102.1 |
| | NKG50 | *Bacillus velezensis* SRCM102747 | 99.6 | 96 | CP028211.1 |
| | HFG7 | *Bacillus amyloliquefaciens* DM09 | 98.8 | 95 | JX014631.1 |
| | HFG36 | *Bacillus subtilis* subsp. *subtilis* NCD-2 | 98.2 | 99 | CP023755.1 |
| | HKG21 | *Bacillus safensis* BRM1 | 99.3 | 100 | CP018100.1 |
| | HKG43 | *Bacillus velezensis* SRCM102747 | 98.9 | 96 | CP028211.1 |
| | HKG45 | *Bacillus velezensis* SRCM102747 | 99.2 | 94 | CP028211.1 |
| | NFG1 | *Bacillus velezensis* P34 | 99.2 | 99 | CP040378.1 |
| | RFG14 | *Bacillus velezensis* P34 | 99.4 | 98 | CP040378.1 |
| | RKG29 | *Bacillus halotolerans* MBH1 | 99.4 | 100 | CP070976.1 |
