## Supplementary material for "Endophytic bacteria *Bacillus velezensis* NKG50 as a potential biocontrol agent of Ascochyta blight on chickpea": Table S2

**Table S2. Genome-to-genome comparison analyses against *B. velezensis* NKG50 sequence.** ANI, average nucleotide identity; dDDH, digital DNA-DNA hybridization. In bold, *B. velezensis* co-species.

| Genomes references | ANI% | dDDH | Genbak accession | Reference |
|---|---|---|---|---|
| ***Bacillus amyloliquefaciens*** subsp. ***plantarum* FZB42** | **98.8** | **91.4** | CP000560.1 | Chen et al., 2007 |
| ***Bacillus velezensis* KACC 13105** | **98.0** | **84.3** | JTKJ00000000.2 | Dunlap et al., 2016 |
| ***Bacillus velezensis* NRRL B-41580T** | **98.0** | **84.5** | NZ_LLZC00000000.1 | Dunlap et al., 2016 |
| ***Bacillus velezensis* JS25R** | **98.0** | **84.6** | NZ_CP009679.1 | NCBI |
| ***Bacillus velezensis* CAU B946** | **97.4** | **79.2** | NC_016784.1 | Blom et al., 2012 |
| *Bacillus siamensis* KCTC 13613T | 94.2 | 56.7 | NZ_AJVF00000000.1 | Jeong et al., 2012 |
| *Bacillus amyloliquefaciens* DSM7 | 93.8 | 55.4 | FN597644.1 | Borriss et al., 2011 |
| *Bacillus subtilis* subsp. *subtilis* 168 | 81.2 | 20.9 | AL009126.3 | Kunst et al., 1997 |
| *Bacillus subtilis* subsp. *spizizenii* TU-B-10 | 81.2 | 21.0 | CP002905.1 | Earl et al., 2012 |
| *Bacillus tequilensis* KCTC 13622T | 81.1 | 70.0 | NZ_AYTO00000000.1 | NCBI |
| *Bacillus mojavensis* UCMB5075 | 80.9 | 21.0 | CP051464.1 | NCBI |
| *Bacillus subtilis inaquosorum* KCTC 13429T | 80.9 | 20.9 | CP029465.1 | NCBI |
| *Bacillus atrophaeus* 1942 | 80.8 | 21.0 | CP002207.1 | Gibbons et al., 2011 |
| *Bacillus licheniformis* ATCC 14580 | 78.7 | 19.6 | NC_006322.1 | Veith et al., 2004 |
| *Bacillus altitudinis* GR-8 | < 70.0 | 19.2 | CP009108.1 | NCBI |
| *Bacillus pumilus* SAFR-032 | < 70.0 | 19.0 | CP000813.4 | NCBI |
| *Bacillus safensis* FO-36b | < 70.0 | 18.5 | ASJD00000000.1 | NCBI |
| *Bacillus xiamenensis* HYC-10 | < 70.0 | 18.0 | AMSH00000000.1 | Lai et al., 2012 |
